# Binding Free energy Decomposition and Multiple Unbinding Paths of Buried Ligands in a PreQ_1_ Riboswitch

**DOI:** 10.1101/2021.07.13.452201

**Authors:** Guodong Hu, Huan-Xiang Zhou

## Abstract

Riboswitches are naturally occurring RNA elements that control bacterial gene expression by binding to specific small molecules. They serve as important models for RNA-small molecule recognition and have also become a novel class of targets for developing antibiotics. Here, we carried out conventional and enhanced-sampling molecular dynamics (MD) simulations, totaling 141.5 μs, to characterize the determinants of binding free energies and unbinding paths for the cognate and synthetic ligands of a PreQ_1_ riboswitch. Binding free energy analysis showed that two triplets of nucleotides U6-C15-A29 and G5-G11-C16, contribute the most to the binding of the cognate ligands, by hydrogen bonding and by base stacking, respectively. Mg^2+^ ions are essential in stabilizing the binding pocket. For the synthetic ligands, the hydrogen-bonding contributions of the U6-C15-A29 triplet are significantly compromised, and the bound state resembles the apo state in several respects, including the disengagement of the C15-A14-A13 and A32-G33 base stacks. The bulkier synthetic ligands lead to significantly loosening of the binding pocket, including extrusion of the C15 nucleobase and a widening of the C15-C30 groove. Enhanced-sampling simulations further revealed that the cognate and synthetic ligands unbind in almost opposite directions. Our work offers new insight for designing riboswitch ligands.

## INTRODUCTION

Noncoding RNAs mediate essential cellular processes such as gene expression and their dysregulation is linked to infectious diseases and cancer (1,2). They can fold into intricate three-dimensional structures with pockets that potentially serve as binding sites for small molecules (3,4). There is growing interest in developing RNA-binding small molecules as therapeutics and chemical probes (5,6). Riboswitches are structured non-coding RNA elements that occur in 5’ untranslated regions of mRNA, most often in bacteria (7). A riboswitch typically consists of two domains: a conserved aptamer domain that folds into a structure with a binding pocket for a ligand molecule, and an expression platform that interfaces with the transcriptional or translational machinery. Directed by the presence or absence of the ligand, the two domains compete for a switch sequence, resulting in two alternative structures for the expression platform that correspond to the on and off states of the mRNA. Ligands range from nucleobases, cofactors, and amino acids to metal ions (8–15). Riboswitches bind their cognate ligands with high affinity and high selectivity. These important properties make riboswitches prime targets for developing small-molecule antibiotics and chemical tools (16–23).

The smallest known aptamer domain is from the class 1 PreQ_1_ riboswitch. With 33 nucleotides, this aptamer forms a compact H-type pseudoknot when bound with PreQ_1_ (24,25) (Figure 1A). The structure consists of two stems: A-form stem S1 formed by pairing the C1 to G5 bases with the G20 to C16 bases, and pseudoknotted stem S2 with canonical C30-G11 and G33-C9 pairs flanking noncanonical A31-G8 and A32-A10 pairs. The intervening sequences are called loops L1 (U6 to C7), L2 (U12 to C15), and L3 (U21 to A29). Prequeuosine0 (PreQ_0_, or Q_0_ for short) and prequeuosine1 (PreQ_1_, or Q_1_ for short) are precursors (Figure 1B) to the modified guanine nucleotide queuosine. PreQ_1_ and PreQ_0_ have a modest difference in binding affinity (with *K*_D_ at 2.05 ± 0.29 nM and 35.10 ± 6.07 nM, respectively) for the aptamer from *Thermoanaerobacter tengcongensis* (*Tte*) (24). Their crystal structures show very similar poses (24–26). PreQ_1_ forms in-plane hydrogen bonds with U6, C15, and A29, and stacks against G5 and C16 on one side and against G11 on the other side (Figure 1A). In the crystal structure of the apo form, the *Tte* aptamer assumes the same fold, but loop L2 in particular is reorganized, with the A14 base inserted into the PreQ_1_-binding pocket whereas the C15 base extruded from the core. Also worth noting are several Mn^2+^ ions (mimicking Mg^2+^) resolved in the latest crystal structure of the PreQ_1_-bound (or apo) *Tte* aptamer (25). The structures of the *Tte* aptamer bound with three synthetic ligands (L_1_ to L_3_; Figure 1C), but with the A13 and A14 (or even C15) nucleobases removed, have been determined (23). The synthetic ligands are bulkier than PreQ_1_, and their binding affinities for the *Tte* aptamer were 50 to 300-fold lower. In the crystal structures, the synthetic ligands, like PreQ_1_ and PreQ_0_, are sandwiched between G5 and C16 on one side and G11 on the other side, but within the ligand plane, only A29 forms a hydrogen bond with the ligands. The reduced number of hydrogen bonds may explain the weakened affinities, but the removal of the L2 nucleobases in the crystal structures complicates the interpretation.

**Figure 1.**
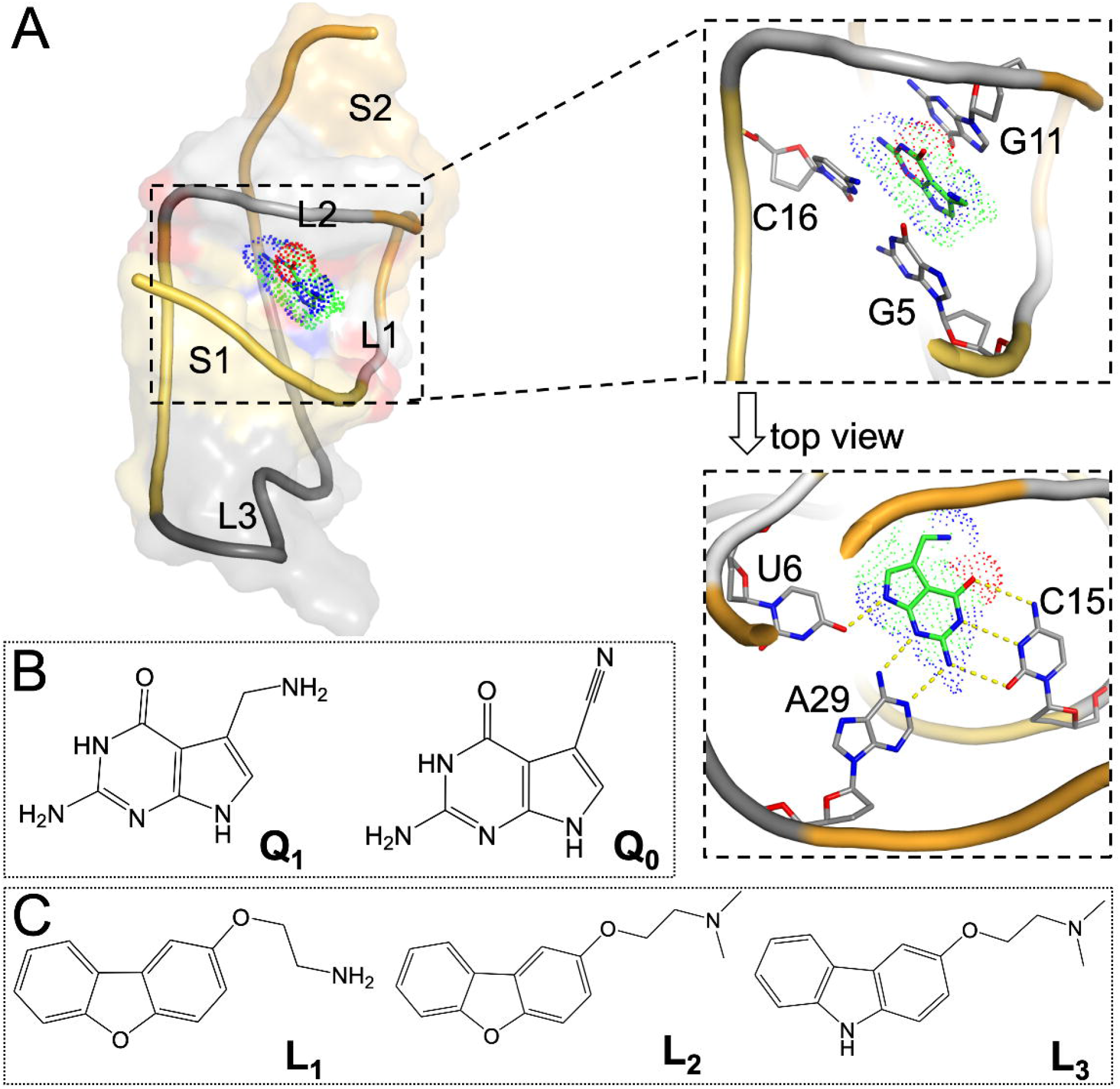
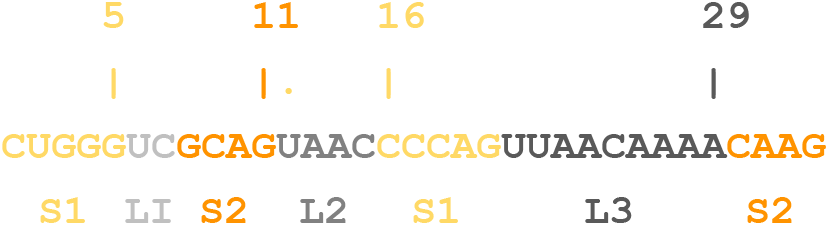
Structures of the PreQ_1_-bound aptamer domain from the *Tte* PreQ_1_ riboswitch and of the cognate and synthetic ligands. (A) Structure of the PreQ_1_-aptamer complex. The aptamer is shown in both cartoon and surface representations, with the following coloring scheme: A zoomed version showing PreQ_1_ in an oblique view is displayed at the right to highlight the base stacking with G11 at the top and with G5 and C16 at the bottom. Below that is a top view highlighting the in-plane hydrogen bonding with U6, C15, and A29. This structure was prepared using coordinates from the PDB entry 6E1W (23), with missing nucleotides copied from PDB entry 3Q50 (24)). (B) Chemical structures of PreQ_1_ and PreQ_0_. (C) Chemical structures of three synthetic ligands, L_1_ to L_3_.

More importantly, crystal structures provide only a single snapshot from an ensemble of conformations. Additional information, in particular energetic and dynamic properties, can come from molecular dynamics (MD) simulations. For example, MD simulations have been used to investigate ligand-induced conformational changes of the aptamer domains from the guanine, adenine, and S-adenosylmethionine sensing riboswitches (27–30). The small size of the PreQ_1_ riboswitch aptamer makes it attractive for MD simulations (31–33). Questions regarding binding affinity and selectivity can be addressed by binding free energy calculations, such as by the molecular mechanics Poisson-Boltzmann surface area (MM-PBSA) method (34). A detailed description of the ligand binding and unbinding paths provide additional insight. As ligand entrance to and exit from a buried site, as found in the PreQ1 riboswitch, occur in timescales usually beyond the capability of conventional MD simulations (35), special techniques are required to speed up the process, such as steer MD (36–38) and metadynamics (39,40). Metadynamics works by adding an external, history-dependent bias potential that acts on a selected number of collective variables.

The folding of RNA requires cations to counter the electrostatic repulsion between backbone phosphates (41,42). Mg^2+^, due to its small radius and double charge, can not only directly or indirectly interact with phosphates in the backbone but also enter the core to interact with nucleobases (41,43). In addition to stabilizing RNA structure (44), Mg^2+^ can help mediate molecular recognition (45,46). Because Mg^2+^ has the same electrons as water, locating Mg^2+^ ions in crystal structures can be a challenge (47) and requires relatively high resolution (25). In MD simulations, ions, including Mg^2+^, are often randomly distributed, and therefore most of them reside in the solvent (42,48). A number of computational methods have been developed to predict Mg^2+^ sites, including MIB and IonCom based on structures or sequences for proteins (http://bioinfo.cmu.edu.tw/MIB/ (49); https://zhanglab.ccmb.med.umich.edu/IonCom/ (50)), and MetalionRNA and MCTBI based on a knowledge-based anisotropic potential or Monte Carlo sampling for RNA (http://metalionrna.genesilico.pl/ (51); http://rna.physics.missouri.edu/MCTBI (52–55)).

Here we report MD simulation results on the energetics and dynamics of the *Tte* PreQ_1_ riboswitch aptamer in complex with the cognate ligands PreQ_0_ and PreQ_1_ and the synthetic ligands L_1_, L_2_, and L_3_. Mg^2+^ are found to be essential in stabilizing the binding pocket for the cognate ligands. By comparing and contrasting these two groups of ligands, we learn how the chemical (e.g., number of hydrogen bond donors and acceptors) and physical (e.g., molecular size) features of ligands affect binding affinity and ligand exit paths. In particular, the reduction in the number of hydrogen bond donors and acceptors from five in the cognate group (Figure 1B) to one in the synthetic group (Figure 1C) leads to a dramatic loss in hydrogen bonding with nucleobases. The larger sizes of the synthetic group also lead to significant loosening of the binding pocket, including extrusion of the C15 nucleobase and a widening of the C15-C30 groove. Correspondingly, whereas the preferred exit of the cognate ligands is through the front door between G5 and G11, the preferred exit of the synthetic ligands is the back door between C15 and C30.

## METHODS

### Preparation of molecular systems

Initial structures of the PreQ_1_ riboswitch aptamer bound with Q_1_, L_1_, L_2_, and L_3_ were taken from PDB entries 6E1W, 6E1S, 6E1U, and 6E1V, respectively (23). The missing nucleotides were transplanted from an earlier Q_1_-bound structure in PDB entry 3Q50 (24). The Q_0_-bound complex was generated from the Q_1_-bound complex by substituting the ligands. The apo form was generated by stripping Q_1_ from the Q_1_-bound complex. Missing hydrogen atoms of the aptamer were added by using the Leap module in Amber (56). The structures of the ligands were optimized using the Gaussian 16 program (57) at the HF/6–31G* level. Note that our MD simulations were completed before the release of PDB entries 6VUH and 6VUI containing the apo and Q_1_-bound structures, respectively, in which Mn^2+^ ions were resolved.

Each structure was then solvated in a rectangular periodic box of TIP3P (58) water molecules with a 12 Å buffer. Systems were prepared both without and with Mg^2+^. The primary method for adding Mg^2+^ was MCTBI (52–55), which identified seven sites. Alternatively, we placed 16 Mg^2^ ions randomly in the solvent. Additional Na^+^ ions were added to neutralize the charges of systems with and without Mg^2+^ ions. The force field for RNA and ions was AMBER ff14SB (59). Force-field parameters of the ligands were from the restrained electrostatic potential charges and the general Amber force field (60).

### Conventional MD simulations

All MD simulations were carried out by running the AMBER18 package (56). Each system was minimized by 2500 steps of steepest descent and 2500 steps of conjugate gradient. The system was then heated from 100 K to 300 K over 50 ps and maintained at 300 K for 50 ps under constant volume. Subsequently the simulation was at constant temperature and pressure for 50 ps to adjust the solvent density. Up to this point, harmonic restraints at a force constant of 5 kcal/mol·Å^2^ were imposed on all solute atoms except those on the L2 loop and nucleotides 16 and 33. The last step of equilibration was simulation at constant temperature and pressure for 1 ns, without any restraint. The temperature (300 K) was regulated by the Langevin thermostat (61) and pressure by the (1 atm) was regulated by the Berendsen barostat (62). Covalent bonds involving hydrogen atoms were treated with the SHAKE algorithm (63) to allow for a time step of 2 fs. The particle mesh Ewald method was applied to treat long-range electrostatic interactions (64). Finally, eight replicate simulations were carried out for 1 μs at constant temperature and pressure. Snapshots were saved every 10 ps for later analysis.

### Binding free energy calculations

Binding free energies and the decomposition into contributions of individual nucleotides were obtained by the MM-PBSA method (34). For each snapshot of the simulations, the binding free energy was calculated according to Equation (1), where “Δ” means the difference between the complex and the separated RNA and ligand. Δ*E*_ele_ was from the Coulomb interactions between the RNA and ligand partial charges, and Δ*E*_vdW_ was from the van der Waals interactions between RNA and ligand atoms. Δ*G*_pol_ was calculated by numerically solving the Poisson-Boltzmann equation; Δ*G*_nonpol_ was estimated from the solvent-accessible surface area (SASA) as *γ*SASA + *β* (65); and Δ*S* was obtained from normal mode analysis (66).

From each of the eight replicate simulations, 2500 snapshots were extracted from the second 500 ns, resulting in a total of 20,000 snapshots for the MM-PBSA calculations on each ligand. Decomposition into contributions of individual nucleotides was done only for the enthalpic components (Δ*E*_ele_, Δ*E*_vdW_, Δ*G*_pol_ and Δ*G*_nonpol_).

### Other analyses

Vertical distance in Figure 2B and z in Figure 6B were calculated using tcl scripts in VMD (67). All other distances and hydrogen bond formation were determined by the CPPTRAJ program (68). Hydrogen bond criteria were doner-acceptor distance < 3.5 Å and donor-H-acceptor angle > 120°. Densities of ligands and of Mg^2+^ were calculated from saved snapshots of the cMD simulations by using the *water-hist* program in the LOOS package (69).

**Figure 2.**
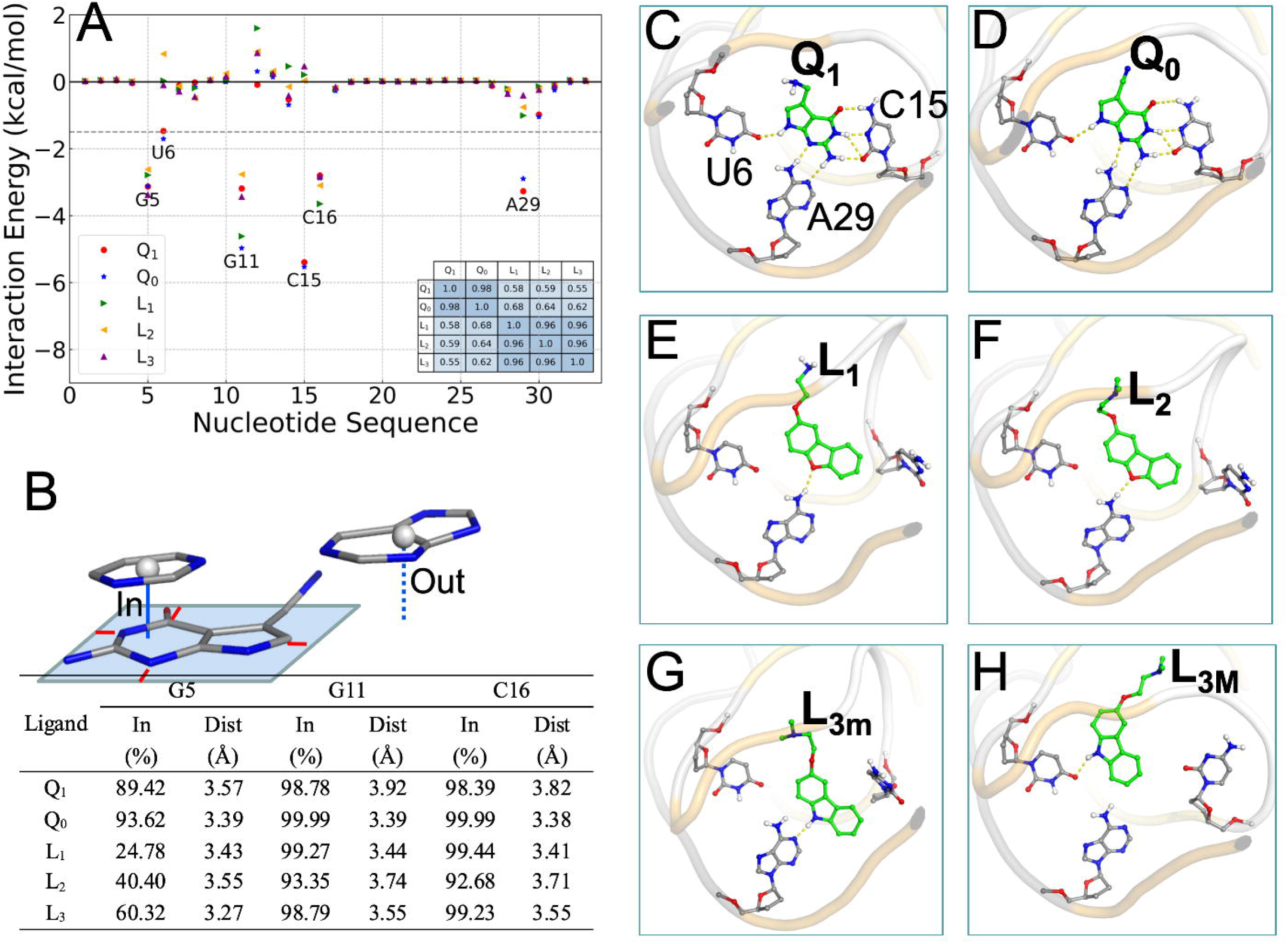
Interactions of cognate and synthetic ligands with the PreQ_1_ aptamer in cMD simulations with Mg^2+^. (A) Contributions of individual nucleotides to the binding free energies. A dashed horizontal line is drawn at −1.5 kcal/mol, which separates the pocket-lining nucleotides from the rest of the sequence. Inset: a table listing the correlation coefficients between the individual contributions of any two complexes. (B) Ligand-nucleotide base stacking statistics. Top: illustration of when a nucleobase is (“in”) or is not (“out”) in a stacking position with the ligand. A rectangle is drawn around the ligand rings atoms, with a minimum of 0.5 Å separation (shown by red lines). A cytosine is in an “in” position, with vertical distance drawn as a solid line, as the projection of its center is inside the rectangle; a guanine is in an “out” position, with vertical distance drawn as a dashed line, as the projection of its center is outside the rectangle. Bottom: in-fractions of three nucleobases and their average vertical distances from the ligand rings. (D)-(H) In-plane hydrogen bonds between ligands and nucleobases, shown as dashed lines, in representative structures from cMD simulations.

### Well-tempered metadynamics simulations

Well-tempered metadynamics simulations were as described (40). In this method, Gaussian functions were applied to fill up wells in the potential of mean force for a collective variable. We chose the distance r from the center of the ligand to the center of the binding pocket (defined by the six nucleotide G5, U6, G11, C15, C16, and C30) as the collective variable. The Gaussian widths was set to 0.5 Å, and the Gaussian hills height was initially set at 1.2 kcal/mol and was gradually decreased with a bias factor of 15 over the course of the simulation. A soft harmonic restraining potential was also applied on the center of the ligand to keep the ligand close to the RNA. Fifteen metadynamics simulations were run for 500 ns each, utilizing the PLUMED v2.3 (70) plugin to the GROMACS 5.1.2 package (71).

## RESULTS

We carried out a total of 141.5 μs MD simulations for the *Tte* PreQ_1_ riboswitch aptamer bound with the cognate ligand Q_1_ or Q_0_, or with the synthetic ligand L_1_, l_2_, or L_3_, or in the apo form (Table 1). Simulations were done with and without Mg^2+^ in order to uncover the effects of this important ion. In addition to conventional MD (cMD), metadynamics simulations were also run to investigate the unbinding paths of ligands. The main results are presented from the simulations with Mg^2+^; only when comparison is made we show Mg^2+^-free results.

**Table 1.** Systems for molecular dynamics (MD) simulations.

| System | # of Mg<sup>2+</sup><br>ions | Type of MD | # of<br>replicates | Length of<br>trajectory(μs) | Total time<br>(μs) |
| --- | --- | --- | --- | --- | --- |
| Apo, Q<sub>1</sub>, Q<sub>0</sub>, L<sub>1</sub>,<br>L<sub>2</sub>, L<sub>3</sub> | 7<sup>a</sup> | cMD | 8 | 1 | 48 |
| Apo, Q<sub>1</sub>, Q<sub>0</sub>, L<sub>1</sub>,<br>L<sub>2</sub>, L<sub>3</sub> | No | cMD | 8 | 1 | 48 |
| Apo | 16<sup>b</sup> | cMD | 8 | 1 | 8 |
| Q<sub>1</sub>, Q<sub>0</sub>, L<sub>1</sub>, L<sub>2</sub>,<br>L<sub>3</sub> | 7<sup>a</sup> | metadynamics | 15 | 0.5 | 37.5 |

<sup>a</sup>Ions initially placed using MCTBI (52-55).
<sup>b</sup>Ions initially placed randomly.

### Free energy decomposition reveals how physicochemical features of ligands affect binding affinity

To explain the significant difference in binding affinity between the cognate and synthetic groups of ligands, we calculated the binding free energies by applying the MM-PBSA method to the second half of eight replicate cMD simulations of each RNA-ligand complex. MM-PBSA has been successfully used on many RNA-ligand (29,30) and protein-ligand (72,73) systems. Although this method can have large uncertainties in the absolute free energy calculated for a given ligand, the relative difference in binding free energy between ligands calculated from comparative MD simulations can be very informative. The binding free energy consists of five terms:

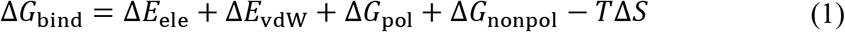

where Δ*E*_ele_ represents the average electrostatic interaction energy between the RNA and the ligand in gas phase; Δ*E*_vdW_ is the counterpart for van der Waals interactions; Δ*G*_pol_ and Δ*G*_nonpol_ account for the solvent environment of the RNA-ligand complex; Δ*S* is the change in conformational entropy upon binding; and *T* is the absolute temperature. These components and the total binding free energies for the cognate and synthetic ligands, calculated from the simulations in the presence of Mg^2+^, are listed in Table 2.

**Table 2.**
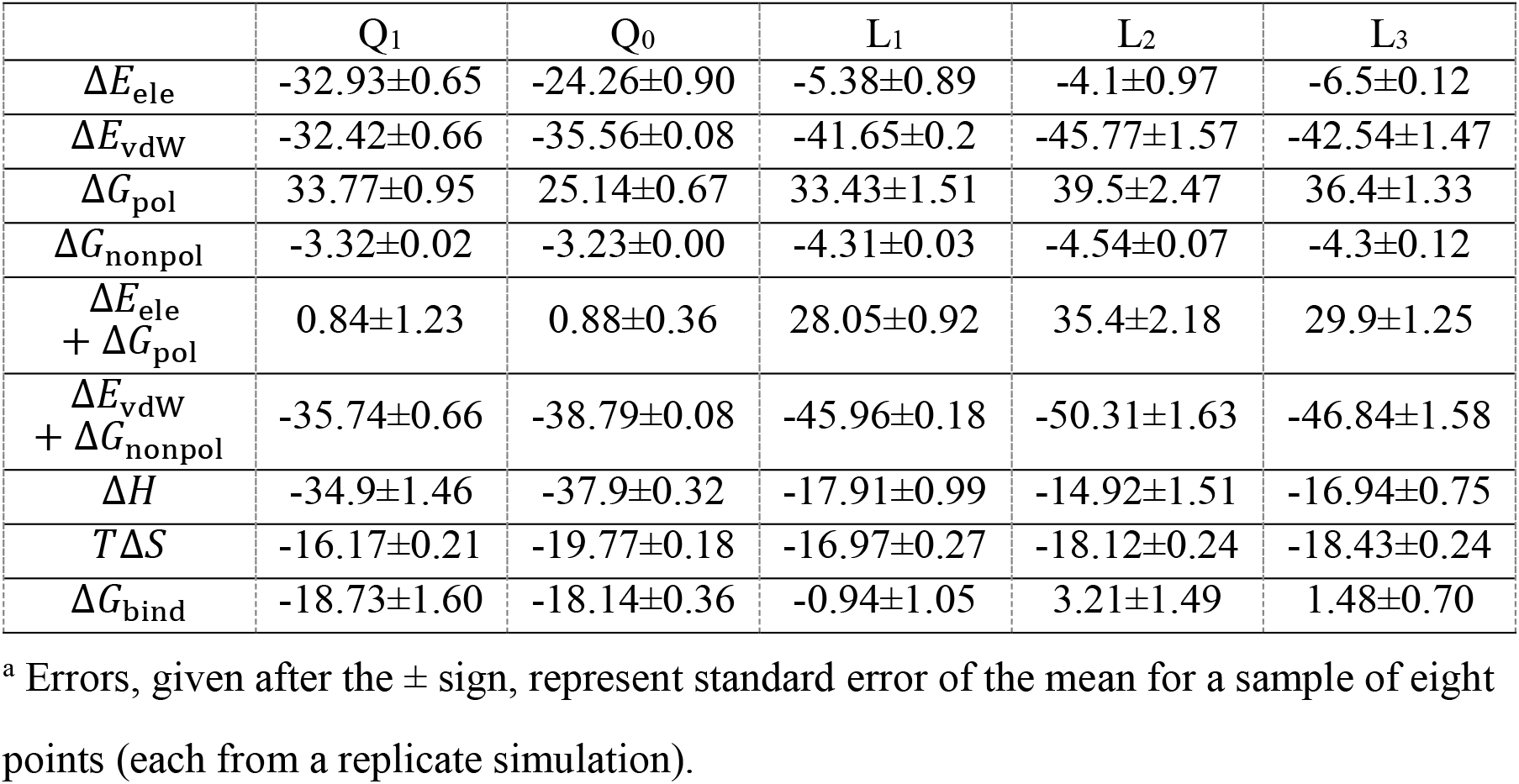
Binding free energies and their components (in kcal/mol) for five ligands.^a^

|  | Q<sub>1</sub> | Q<sub>0</sub> | L<sub>1</sub> | L<sub>2</sub> | L<sub>3</sub> |
| --- | --- | --- | --- | --- | --- |
| $\Delta E_{\text{ele}}$ | -32.93±0.65 | -24.26±0.90 | -5.38±0.89 | -4.1±0.97 | -6.5±0.12 |
| $\Delta E_{\text{vdW}}$ | -32.42±0.66 | -35.56±0.08 | -41.65±0.2 | -45.77±1.57 | -42.54±1.47 |
| $\Delta G_{\text{pol}}$ | 33.77±0.95 | 25.14±0.67 | 33.43±1.51 | 39.5±2.47 | 36.4±1.33 |
| $\Delta G_{\text{nonpol}}$ | -3.32±0.02 | -3.23±0.00 | -4.31±0.03 | -4.54±0.07 | -4.3±0.12 |
| $\Delta E_{\text{ele}} + \Delta G_{\text{pol}}$ | 0.84±1.23 | 0.88±0.36 | 28.05±0.92 | 35.4±2.18 | 29.9±1.25 |
| $\Delta E_{\text{vdW}} + \Delta G_{\text{nonpol}}$ | -35.74±0.66 | -38.79±0.08 | -45.96±0.18 | -50.31±1.63 | -46.84±1.58 |
| $\Delta H$ | -34.9±1.46 | -37.9±0.32 | -17.91±0.99 | -14.92±1.51 | -16.94±0.75 |
| $T\Delta S$ | -16.17±0.21 | -19.77±0.18 | -16.97±0.27 | -18.12±0.24 | -18.43±0.24 |
| $\Delta G_{\text{bind}}$ | -18.73±1.60 | -18.14±0.36 | -0.94±1.05 | 3.21±1.49 | 1.48±0.70 |

<sup>a</sup> Errors, given after the ± sign, represent standard error of the mean for a sample of eight points (each from a replicate simulation).

MM-PBSA predicted binding free energies of ~ −18 kcal/mol for the cognate group and close to ~1 kcal/mol for the synthetic group. Though these results exaggerates the difference in Δ*G*_bind_ (the experimental difference is ~2 kcal/mol (23,24)), they do correctly predict the cognate group as the stronger binders. Comparing the two groups of ligands, the polar components (Δ*E*_ele_ + Δ*G*_pol_) are much more favorable (by ~30 kcal/mol) to the cognate group, offset only partially (by ~10 kcal/mol) by the nonpolar components (Δ*E*_vdW_ + Δ*G*_nonpol_) that favors the synthetic group. These contrasts can be easily attributed to the greater number of hydrogen bond doners and acceptors in the cognate group and the bulkier sizes of the synthetic group.

To gain insight into how RNA-ligand interactions lead to the difference in affinity between the cognate and synthetic groups, we decomposed Δ*G*_bind_ into contributions of the 33 nucleotides of the *Tte* aptamer (Figure 2A). The correlation coefficients of these individual contributions for any two ligands are nearly 1 within both the cognate and synthetic groups, but reduces to ~0.6 between the groups (Figure 2A, inset table). This correlation analysis clearly indicates that the two groups of ligands are distinct. The only nucleotides that contribute more than 1.5 kcal/mol in at least one complex are the six that form either base stacking (G5, G11, C16) or in-plane hydrogen bonding (U6, C15, A29) with the ligands (Figure 1A). The base-stacking nucleotides contribute nearly the same to the binding free energies of the two groups of ligands, but the hydrogen-bonding nucleotides differ by 1.8, 5.7, and 2.4 kcal/mol, respectively, in their contributions to the two groups. This result identifies in-plane hydrogen bonding as the dominant factor for the difference in affinity between the cognate and synthetic groups.

The MM-PBSA results calculated from the simulations in the absence of Mg^2+^ are presented in Table S1 and Figure S1A. The qualitative differences between the cognate and synthetic ligands described above are also valid in the Mg^2+^-free simulations. The only major change is that the binding free energies of the cognate group become less favorable by ~10 kcal/mol, which come entirely from the polar components. The stabilization of cognate ligand binding by Mg^2+^ is explained below.

### Binding poses of the five ligands differ in overt and subtle ways

Next we present structural differences around the binding pocket among the aptamer-ligand complexes in the cMD simulations. To characterize base stacking, we calculated the fraction (“in-fraction”) of snapshots where a nucleobase falls within a rectangle around the ring atoms of the ligand, and among these snapshots, the vertical distance between the center of the nucleobase and the rectangle (Figure 2B, top). For the cognate ligands, the in-fractions of G5, G11, and C16 all are close to 100%, but for the synthetic ligands, the in-fractions of G5 decrease to between 25% to 60% (Figure 2B, bottom). Among the in-fractions, the distances between the nucleobases and the ligand rings are about 3.5 Å, the van der Waals contact distance for carbon atoms. A subtle but consistent difference within the cognate group is that all the three nucleobases have slightly higher in-fractions and shorter distances, thus implicating a tighter binding pocket, for Q_0_ than for Q_1_. The tighter binding pocket for Q_0_ is consistent with the more favorable van der Waals interaction energy and the higher entropic cost listed in Table 2. In the absence of Mg^2+^, the in-fractions of G5 decrease to below 50% for the cognate group (Figure S1B). The effect of Mg^2+^ on base stacking is more subtle for the synthetic group, with the in-fractions of G11 and C16 decreasing to between 71% to 85%.

The three nucleobases, U6, C15, A29, form six in-plane hydrogen bonds with the cognate ligands in the crystal structures (Figure 1A) (23–26). These hydrogen bonds are well maintained in the cMD simulations with Mg^2+^ (Table 3). Figure 2C,D shows the typical poses of Q_1_ and Q_0_, respectively, in the binding pocket. In particular, the C15 nucleobase remains parallel to the ligand rings and forms three stable hydrogen bonds with Q_1_ and Q_0_, pairing N4, N3, and O2 of C15 with O1, N1, and N5 of Q_1_ and Q_0_. About one third of the time, O2 of C15 and N1 of Q_1_ and Q_0_ form an additional hydrogen bond. In contrast, the rings of the synthetic ligands have a single hydrogen bond donor or acceptor (compared with five for the cognate ligands) and forms a single hydrogen (Table 3). For L_1_ and L_2_, the O2 acceptor pairs with the A29 N6 donor, and the C15 nucleobase extrudes into an orthogonal orientation, to accommodate the synthetic ligands’ bulkier size (Figure 2E,F). In the crystal structures of the aptamer bound with the synthetic ligands (23), L_3_ is positioned similarly to L_1_ and L_2_ and also hydrogen bonds with A29. However, unlike L_1_ and L_2_, L_3_ is a donor, not acceptor, and correspondingly the partner changes from N6 to N1 of A29. As a result of this change in hydrogen bonding partner, the rings of L_3_ move closer to C30 (Figure 2G). In the cMD simulations, hydrogen bonding with A29 is found in only 27.6% of the snapshots (Table 3). Instead, in 55.2% of the snapshots, L_3_ moves laterally to hydrogen bond with another acceptor, i.e., O4 of U6, allowing the C15 nucleobase to keep its parallel orientation (Figure 2H). We label the A29-hydrogen bonded minor and the U6-hydrogen bonded major poses as L_3m_ and L_3M_, respectively. The L_3m_ and L_3M_ poses readily interconvert, with multiple transitions observed in each simulation (Figure S2).

**Table 3.** Hydrogen bonding probabilities.

| Donor | Acceptor | Probability (%) |  |
| --- | --- | --- | --- |
|  |  | w/ Mg<sup>2+</sup> | w/o Mg<sup>2+</sup> |
| Q<sub>1</sub> / Q<sub>0</sub> N4 | U6 O4 | 85.4 / 95.4<sup>a</sup> | 92.1 / 95.6 |
| C15 N4 | Q<sub>1</sub> / Q<sub>0</sub> O1 | 94.9 / 92.2 | 42.6 / 18.9 |
| Q<sub>1</sub> / Q<sub>0</sub> N1 | C15 N3 | 91.3 / 95.2 | 43.7 / 19.4 |
| Q<sub>1</sub> / Q<sub>0</sub> N5 | C15 O2 | 87.7 / 96.4 | 52.2 / 69.0 |
| Q<sub>1</sub> / Q<sub>0</sub> N1 | C15 O2 | 30.7 / 45.1 | 71.1 / 87.8 |
| A29 N6 | Q<sub>1</sub> / Q<sub>0</sub> N2 | 87.8 / 98.5 | 98.8 / 99.7 |
| Q<sub>1</sub> / Q<sub>0</sub> N5 | A29 N1 | 85.5 / 82.6 | 95.8 / 93.8 |
| *Q<sub>1</sub> N3* | *G5 O6 / N7* | *21.6 / 18.9<sup>b</sup>* | *50.3 / 6.4* |
| *Q<sub>1</sub> N3* | *G11 N7* | *32.6* | *20.1* |
| A29 N6 | L<sub>1</sub> / L<sub>2</sub> O2 | 86.5 / 76.7 | 76.4 / 94.0 |
| L<sub>3</sub> N1 | U6 O4 / A29 N1 | 55.2 / 27.6 | 52.9 / 14.0 |
| *L<sub>1</sub> N1* | *G5 O6 / N7* | *37.0 / 52.3* | *7.8 / 16.9* |

<sup>a</sup> “/” separates the hydrogen bonding probabilities for two different donors or acceptors listed in the same row.
<sup>b</sup> Entries listed in italic are for Q<sub>1</sub> and L<sub>1</sub> methylamines as hydrogen bond donors.

Q_1_ differs from Q_0_ by the substitution of a methylamine for a cyano (Figure 1B). In 40.5% of the snapshots, this methylamine hydrogen bonds with O6 or N7 of G5; in another 32.6% of the snapshots the hydrogen bond partner switches to N7 of G11 (Table 3). Thus the methylamine group of Q_1_, by changing the C3-C5-C6-N3 torsion angle, alternates its hydrogen-bonding partner between G5 and G11. In the recent crystal structure [Protein Data Bank (PDB) entry 6VUI] (25), the methylamine group hydrogen bonds with G5, though the authors did consider but dismissed G11 as an alternative partner. L_1_ also has a methylamine, and it too can hydrogen bond with O6 or N7 of G5 (Table 3). However, the methylamine in L_1_ is more separated from nearest ring than in Q_1_ and this greater separation prevents hydrogen bonding with G11. Indeed, when the L_1_ methylamine hydrogen bonds with G5, the rings are removed from G5 and this explains why the in-fraction of G5 for L_1_ is only about one half of that for L_2_ (Figure 2B).

Mg^2+^ significantly affects the hydrogen bonding of the cognate ligands with C15 (Table 3; Figures 2C,D and S1C,D). Whereas C15 and Q_1_ / Q_0_ form three stable hydrogen bonds in the simulations with Mg^2+^, only one stable hydrogen bond is formed in the absence Mg^2+^, between C15 O2 and Q_1_ / Q_0_ N1 (or N5). This happens as the C15 nucleobase moves and tilts away from the ligand rings (see below). Since the synthetic ligands do not hydrogen bond with C15, their in-plane hydrogen bonding is not affected by Mg^2+^ (Table 3; Figures 2E-H and S1E-H).

### How does Mg^2+^ stabilize aptamer binding of cognate ligands?

All our MD simulations were carried out before the release of the recent crystal structures of the *Tte* aptamer in apo form and bound with Q_1_ (PDB entries 6VUH and 6VUI) (25), in which three and two Mn^2+^ ions, respectively, were located in the major groove lining the ligand binding pocket. These crystal metal sites thus allowed us to test the MD simulations, with seven Mg^2+^ ions initially placed using MCTBI (52–55) (Figure S3A). As shown in Figures 3A and S3B, the crystal metal sites overlap well with the Mg^2+^ densities calculated in our cMD simulations of the Q_1_-bound and apo forms. The Mg^2+^ densities in the Q_0_-bound form also overlap well with the Q_1_ crystal metal sites (Figure S3C), whereas the densities in the complexes with the synthetic ligands overlap well with the apo crystal metal sites (Figure S3D). We label these sites as M1, M2 (M2’), M3, and M4. In comparison, random initial placement of 16 Mg^2+^ ions was not effective.

**Figure 3.**
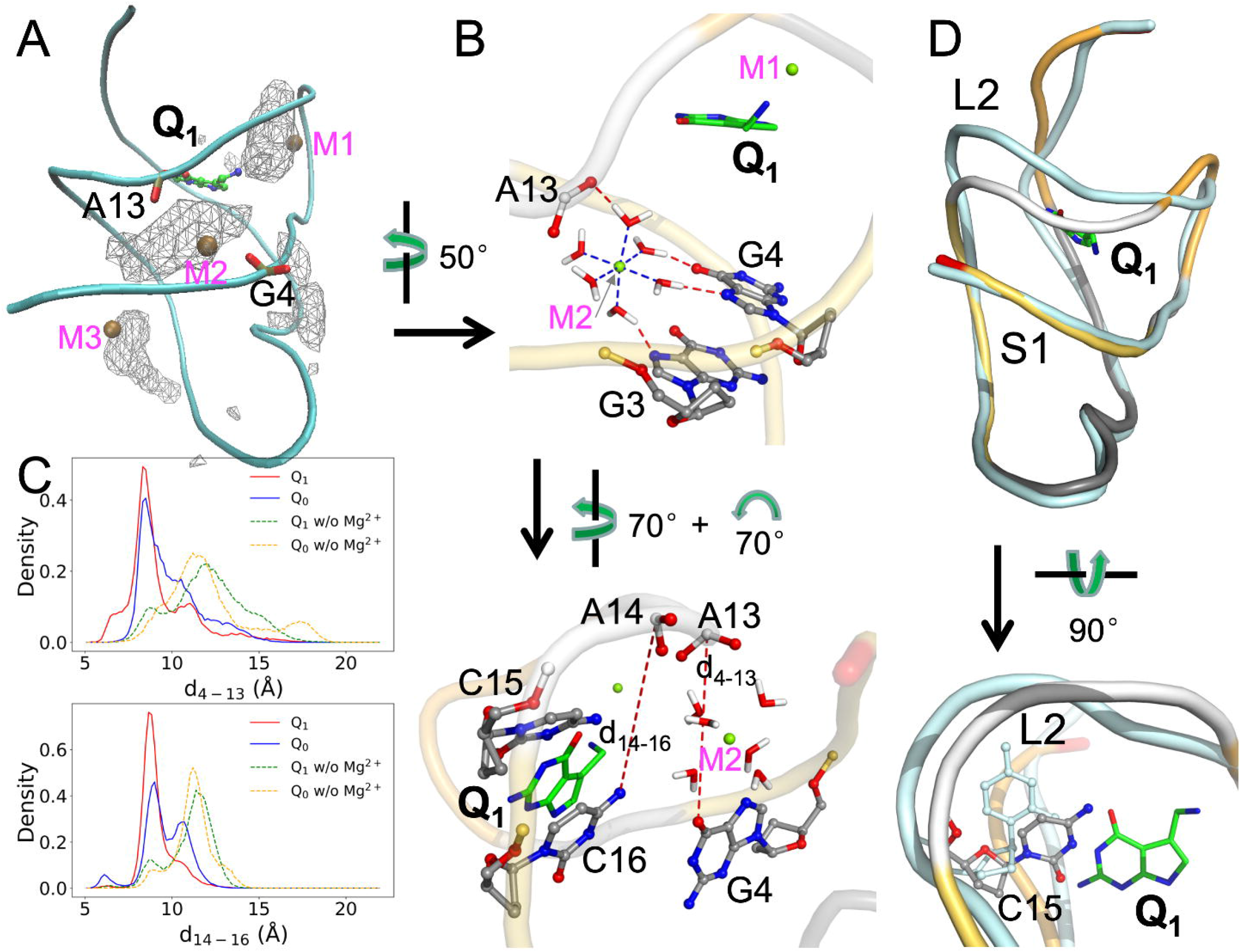
Distributions and effects of Mg^2+^. (A) Density contours of Mg^2+^ ions in the Q_1_-bound complex, shown as wireframe. Three Mn^2+^ ions in PDB entry 6VUI are shown as ochre spheres; the corresponding Mg^2+^ sites are labeled as M1, M2, and M3. Phosphate groups in G4 and A13 are shown in stick representation, to highlight the bridging role of M2. (B) Two views showing the coordination of the Mg^2+^ ion at the M2 site. Top: coordination of Mg^2+^ by six water molecules and the latter’s hydrogen bonding with the G3 and G4 nucleobases and A14 phosphate. Bottom: two distances, d_4-13_ (between A4 O6 and A13 OP1 and d_14-16_ (between A14 P and C16 N4), introduced to characterize the effects of the Mg^2+^ ion. (C) Probability densities of d_4-13_ and d_14-16_, in cMD simulations with and without Mg^2+^ ions. (D) Effect of Mg^2+^ ions on the separation of the L2 loop from the S1 helix in the Q_1_-bound form. Two representative structures are superimposed, with the aptamer in the presence of Mg^2+^ shown in the same multi-color scheme as in Figure 1A and the aptamer in the absence of Mg^2+^ shown in a uniform cyan color. In the bottom view, the C15 nucleotides in the two structures are shown in a stick representation.

Of particular importance is a deep site, M2, in the complexes with the cognate ligands, where Mg^2+^ bridges between G4 and G3 on the S1 helix and A13 on the L2 loop (Figures 3B). To present a full picture of this bridging effect in the MD simulations, we monitored two distances between S1 and L2: d_4-13_ between A4 O6 and A13 OP1; and d_14-16_ between A14 P and C16 N4 (Figures 3B, bottom panel). The probability densities of these two distances are both peaked around 8.5 Å (Figures 3C). In the simulations without Mg^2+^, the peaks shift to larger distances by 2 to 4 Å, indicating a greater separation of L2 from S1. The increased separation is also evident when representative structures from the RNA-Q_1_ (or RNA-Q_0_) simulations with and without Mg^2+^ are superimposed (Figure 3D, top panel). The further separation of L2 from S1 creates new room for C15 (Figure 3D, bottom panel), which as noted above moves and tilts away from the cognate ligand rings in the absence of Mg^2+^ (Figures 2C and S1C). Very similar differences are also observed in the simulations of the Q_0_-bound aptamer with and without Mg^2+^ (Figures 3C and S3E; also compare Figures 2D and S1D). Therefore Mg^2+^ is essential in maintaining C15 in a position to form stable hydrogen bonds with the cognate ligands.

### The Aptamer bound with synthetic ligands resembles the apo state

Connolly et al. (23) recognized two key differences between the Q_1_- and L_1_-bound structures and concluded that the latter structure is similar to the apo state. The first, already described above, is the extrusion of the C15 nucleobase into an orthogonal orientation (Figure 2C,E). The second is a farther separation of the A32 and G33 nucleobases from the ligand rings in the L_1_-bound structure. However, the latter crystal structure was determined with the A13 and A14 nucleobases removed, and thus represented an incomplete picture. Our MD simulations of the complete aptamer now provide strong additional support for the conclusion that the structures bound with the synthetic ligands are similar to the apo state.

First, the L2 loop adopts distinct conformations in the apo and Q_1_-bound structures (PDB entries 6VUH and 3Q50 (24,25); Figure 4A, left panel). In the apo structure, L2 is closer toward the ligand binding pocket to allow the insertion of the A14 nucleobase into the pocket. L2 in our simulations of the L_1_-bound aptamer adopts a similar “close” conformation, although the A14 nucleobase is extruded to accommodate the presence of the ligand (Figure 4A, right panel). Second, as noted above, Mg^2+^ densities in our simulations of the Q_1_-bound aptamer overlap with Mn^2+^ ions found in the Q_1_-bound structure (PDB entry 6VUI) (25) (Figure 3A and Figure 4B, left panel). In contrast, Mg^2+^ densities in the L_1_-bound aptamer overlap with Mn^2+^ ions found in the apo structure (PDB entry 6VUH) (25) (Figure S3D and Figure 4B, right panel). Finally and most importantly, in our simulations of the Q_1_-bound aptamer, three nucleotides from the L_2_ loop, C15, A14, and A13, maintain continuous base stack with two nucleotides, A32 and G33 (the Shine-Dalgarno sequence), in the S2 helix (Figure 4C, left panel). The continuous base stack is crucial for inhibiting gene expression by sequestrating the Shine-Dalgarno sequence from recognition by the ribosome (25). However, in the simulations of the L_1_-bound aptamer, the L2 nucleotides and the S2 nucleotides form two separate stacks (Figure 4C, right panel). Both A14 and A13 take the orthogonal orientation of C15 to form a base stack that is disengaged from the A32-G33 stack. In the apo crystal structure, both C15 and A13 have the orthogonal orientation (Figure 4A, left panel).

**Figure 4.**
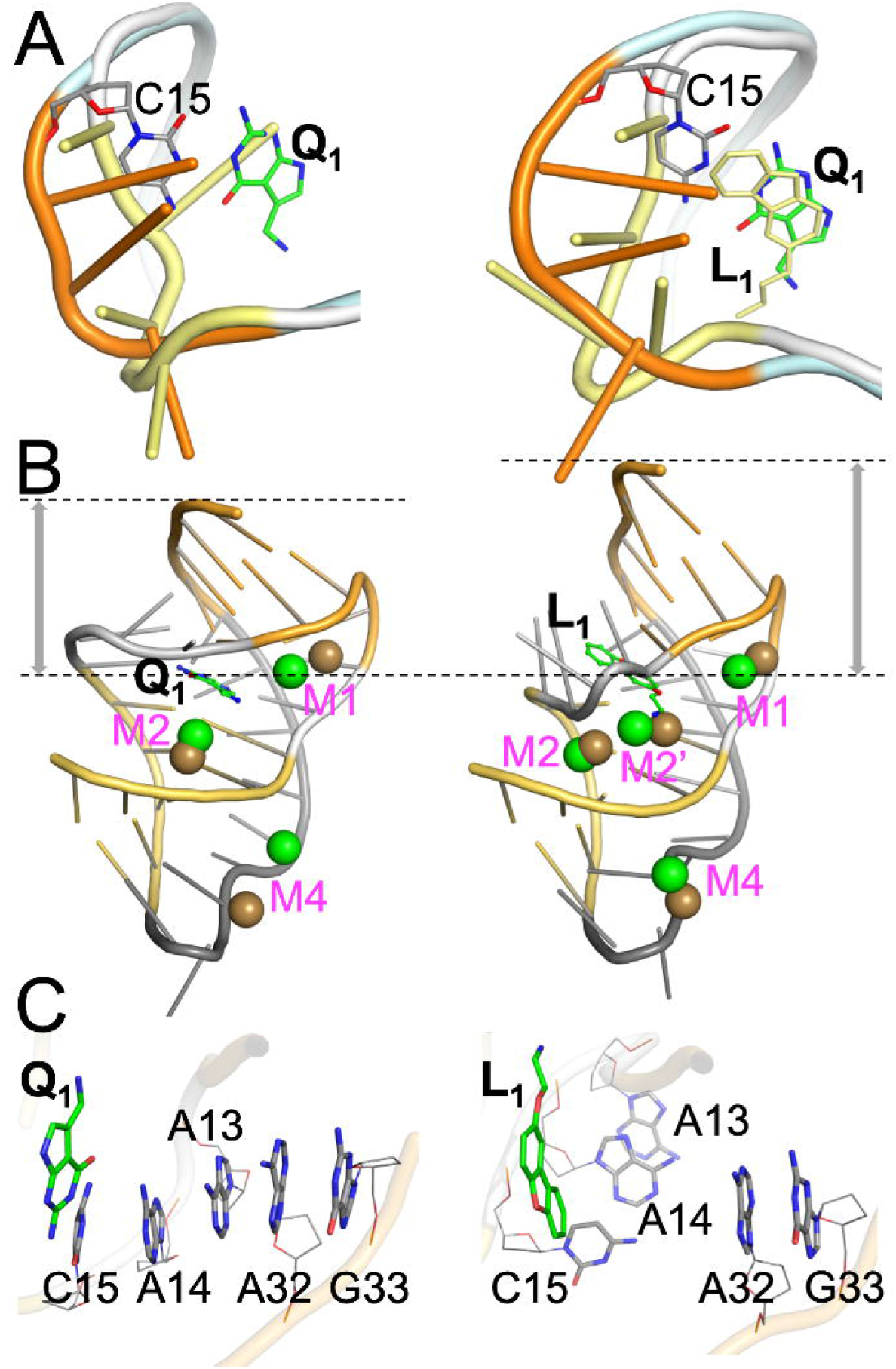
Contrast between Q_1_- and L_1_-bound complexes. (A) Conformations of the L2 loop (U12-A13-A14-C15). Left: conformations adopted in the apo (yellow) and Q_1_-bound (orange) forms, from PDB entries 6VUH and 3Q50, respectively. Right: conformations in representative structures of the L_1_-bound (yellow) and Q_1_-bound (orange) forms from cMD simulations. (B) Comparison between representative Mg^2+^ sites from cMD simulations and Mn^2+^ positions from crystal structures (PDB entries 6VUI and 6VUH). Left: similarity between cMD Mg^2+^ (green) and crystal Mn^2+^ (ochre) positions in the Q_1_-bound form. Right: similarity between cMD Mg^2+^ (green) positions in the L_1_-bound from and crystal Mn^2+^ (ochre) positions in the apo form. The L_1_-bound structure has a larger separation between the ligand and the 3’ end. (C) The orthogonal orientations of the C15-A14-A13 nucleobases in the Q_1_- and L_1_-bound forms in cMD simulations, leading to one stack or two separate stacks, respectively, with the A32-G33 nucleobases.

### The synthetic ligands lead to loosening at the back of the binding pocket

We now examine the total volume explored by the atoms of each ligand in the cMD simulations, by calculating the density contour of the ligand (Figures 5A-C). In line with the foregoing observation that the binding pocket is tight for Q_0_ (Figure 2B, bottom), this cognate ligand shows a very compact density contour (Figure 5A, green). The density contour of Q_1_ (Figure 5A, red) is slightly expanded around the rings, and there is also extra density for the methylamine “head” group.

**Figure 5.**
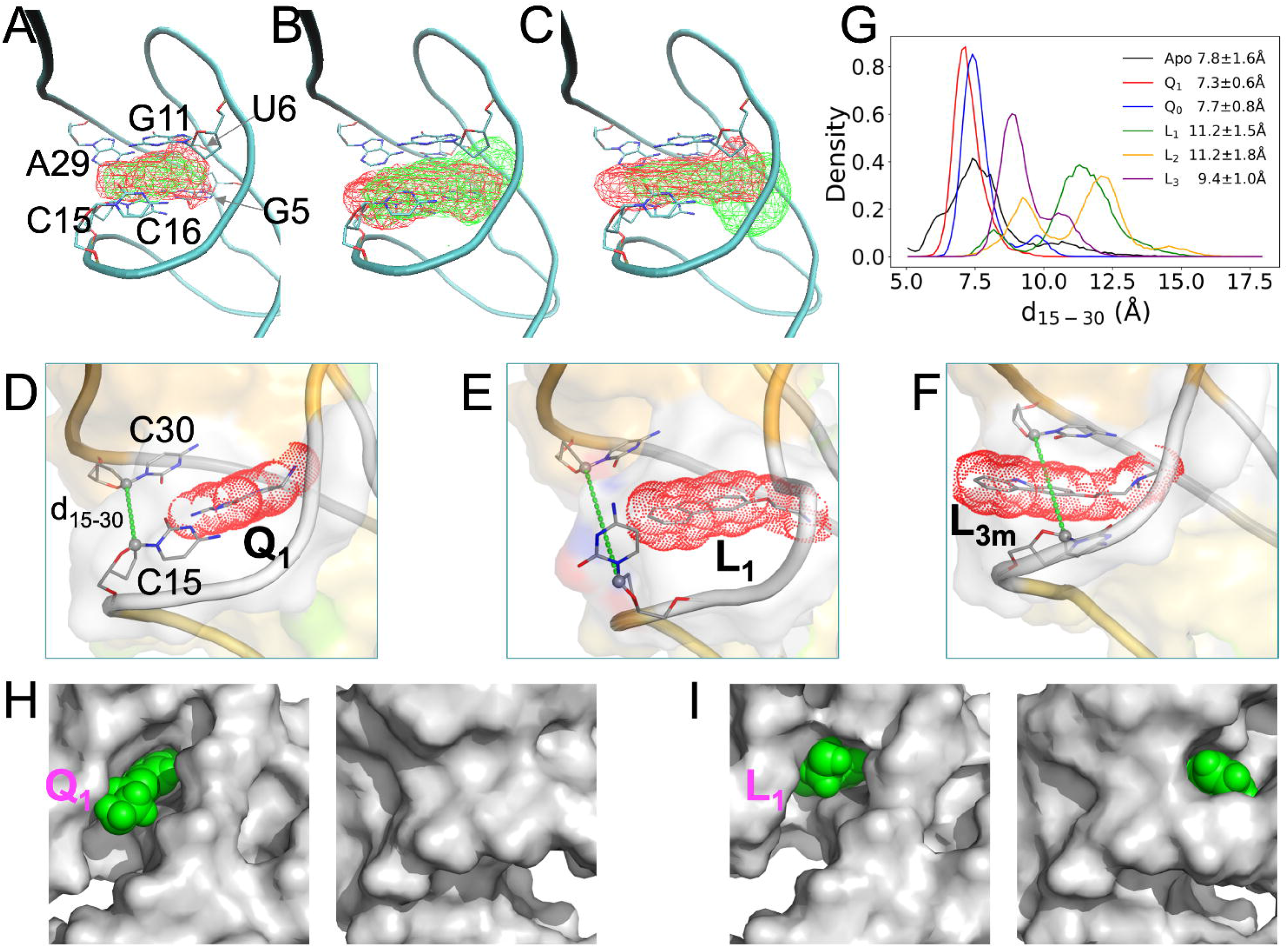
Loosening in the back of the binding pocket when bound with synthetic ligands, in cMD simulations with Mg^2+^. (A)-(C) Density contours of ligands, shown as wireframe in reference to a representative Q_1_-bound structure, with the six pocket-lining nucleotides displayed in stick representation. Panel (A) shows contours in red for Q_1_ and in green for Q_0_; panel (B) shows contours in red for L_1_ and in green for L_2_; and panel (C) shows contours in red for L_3m_ and in green for L_3M_. The L_3m_ and L_3M_ contours were calculated only from snapshots where hydrogen bonding with A29 or U6 was present. (D)-(F) Representative conformations of the Q_1_-, L_1_-, and L_3m_-bound forms, respectively. Ligands are shown in both stick representation and as dot surface. The C1’ atoms of C15 and C30 are connected to define the distance d_15-30_ and to illustrate the back door. (G) The probability densities of d_15-30_ in the apo form and the five liganded forms. (H) Two views into Q_1_ in the binding pocket. The aptamer is shown as gray surface while the ligand is shown as green spheres. Left: front view showing Q_1_ exposure; right: back view showing buried Q_1_. (I) Corresponding presentation for L_1_, except that this ligand is exposed both on the front and on the back.

**Figure 6.**
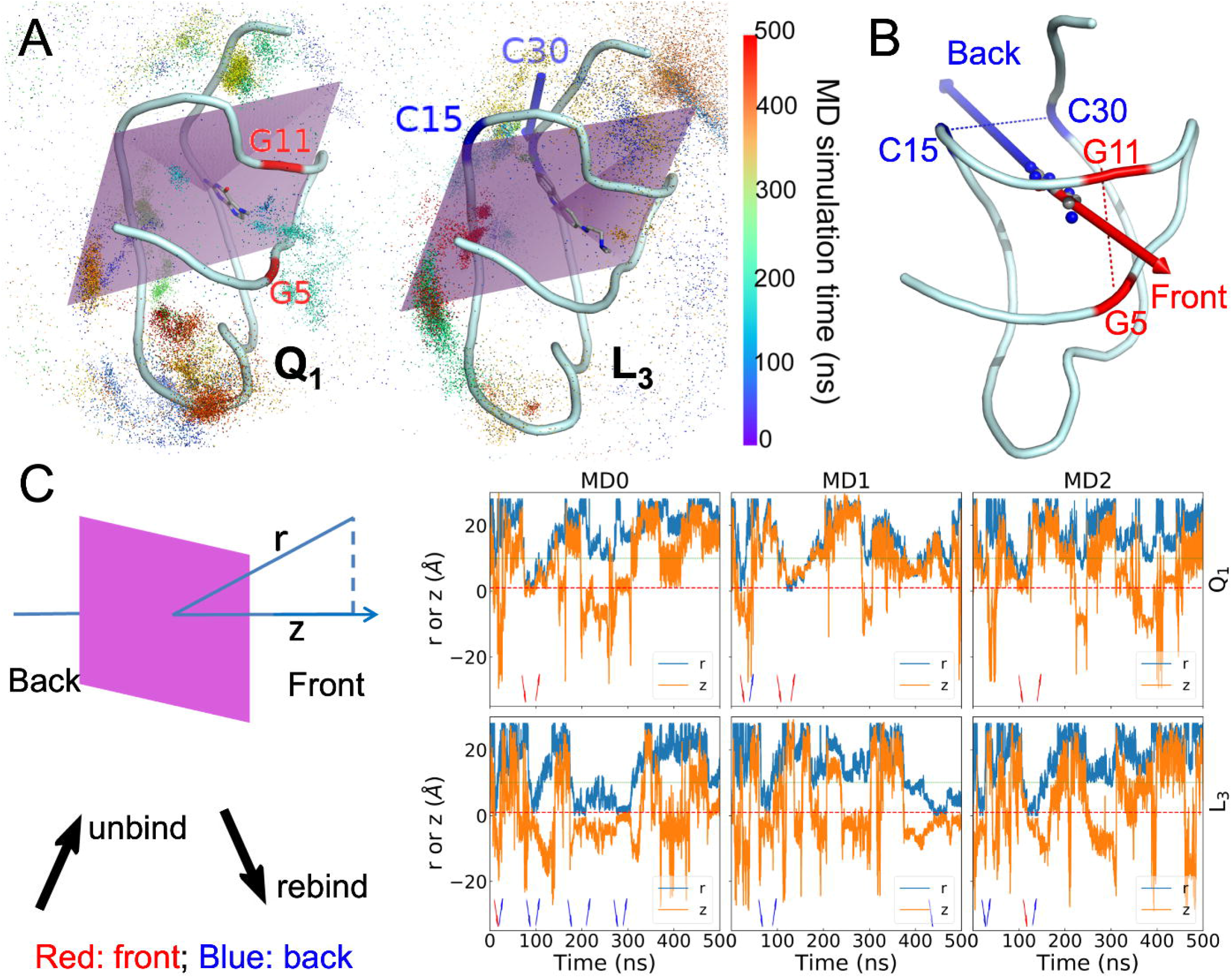
Unbinding and rebinding pathways of ligands. (A) The trajectories of ligand centers shown as dots colored according to the MD simulation time. The aptamer and bound ligands are shown in cartoon and stick representations, respectively. Left: Q_1_; right: L_3_. A plane in purple bisects the binding pocket into the front half and the back half. Two nucleotides defining the front door in the Q_1_-bound complex are labeled in red; two nucleotides defining the back door in the L_3_-bound complex are labeled in blue. (B) The front and back unbinding paths. (C) The center-to-center distance r between the ligand and the binding pocket and the z coordinate of the ligand center along the normal of the pocket-bisecting plane. The left panel illustrates r and z, and presents interpretations of arrow directions and colors that appear in the right three panels, which show time traces of r and z in three metadynamics simulations.

The density contours of L_1_ and L_2_ are further expanded, at both the back (facing the C15-C30 groove) and the front (facing the G5-G11 groove) (Figure 5B). At the back, the expansion, due to the additional ring, would clash with the C15 nucleobase and leads to its extrusion into an orthogonal orientation. At the front, the expansion is largely due to the longer head group (compare Figure 1B and Figure 1C). This front expansion, both longitudinally and laterally, is especially prominent for L_2_, which has two extra terminal methyls in the head group. The aforementioned possibility of the L_1_ methylamine hydrogen bonding with G5 leads to a retraction of the rings toward the back, accounting for the greater back expansion of L_1_ relative to L_2_. As described above, L_3_ readily switches between two poses, L_3m_ (hydrogen bonding with A29) and L_3M_ (hydrogen bonding with U6) (Figure S2). The density contour of L_3m_ (Figure 5C, red) moves slightly farther toward the back than that of L_1_, due to the change in hydrogen bonding partner from N6 to N1 of A29 (Figure 2E, G). Meanwhile, the density contour of L_3M_ (Figure 5C, green) moves toward the front, passing that of L_2_.

The deeper penetration into the back of the binding pocket by L_1_, L_2_, and L_3m_ relative to the cognate ligands are illustrated by snapshots shown in Figures 5D-F and S4A. In one of the eight simulations of the aptamer-L_3_ simulations (Figure S2, MD4), a major portion of the ligand rings is even transiently positioned outside the back door between C15 and C30 (Figure 5F). In two simulations without Mg^2+^, L_3_ escaped altogether through the back door.

To gauge the width of the back door, we monitored the distance (d_15-30_) between the C1’ atoms of C15 and C30 (Figures 5D-F and S4A). The d_15-30_ probability densities in the simulations of the apo and five bound forms are shown in Figure 5G. The mean ± standard deviation of d_15-30_ in the apo form is 7.8 ± 1.6 Å. The mean value is preserved by the cognate ligands (7.3 ± 0.6 Å for Q_1_; 7.7 ± 0.8 Å for Q_0_), but is significantly elevated by the synthetic ligands (11.2 ± 1.5 Å for L_1_; 11.2 ± 1.8 Å for L_2_; 9.4 ± 1.0 Å for L_3_). Similar results are also obtained from the cMD simulations without Mg^2+^ (Figure S4B).

Another distance, d_5-11_, between the Cl’ atoms of G5 and G11 was also monitored to gauge the width of the front door (Figure S4C,D). The five bound forms do not show significant differences in d_5-11_ among themselves, but their mean d_5-11_ values, around 12 Å, are larger by 1 to 2 Å than the apo counterpart. Similar mean d_5-11_ values are found for the five bound forms with and without Mg^2+^, but for the apo form, d_5-11_ shifts to larger values (by ~2 Å) when Mg^2+^ is absent (Figure S4E).

In short, our cMD simulations reveal that, whereas the cognate ligands maintain the intrinsic width of the back door (as found in the apo form), the synthetic ligands widen this door and, in the case of L_3_, can even partially slip through it. In contrast, the front door is kept to approximately the same width by the cognate and synthetic ligands. Consequently, while both the cognate and synthetic ligands are mostly buried in the binding pocket, the cognate ligands are exposed at the front (Figures 5H and S4F) but the synthetic ligands are exposed at both the front and the back (Figures 5I and S4G,H).

### Cognate and synthetic ligands unbind through opposite pathways

The fact that Q_1_ and Q_0_ are exposed only at the front suggests that the cognate ligands bind and unbind through the front door. On the other hand, the exposure of the synthetic ligands at both the front and the back and the (partial) escape of L_3_ through the back door suggest that these ligands may enter and exit through both doors. To investigate the unbinding and rebinding pathways, we carried out 15 well-tempered metadynamics (40) simulations for each. In these simulations, biases were introduced to flatten the potential of mean force along a collective variable, here defined as the distance, r, from the center of the ligand to the center of the binding pocket (lined by the six nucleotides G5, U6, G11, C15, C16, and C30). The biases were gradually reduced during the 500 ns simulations. The trajectories of the ligands are illustrated in Figures 6A and S5A.

To determine whether unbinding or rebinding occurred through the front door (between G5 and G11) or back door (between C15 and C30) (Figure 6B), we introduced a plane passing through the C1’ atoms of U6 and A10 and the C4 atom of C15, which bisects the binding pocket (Figures 6A and S5A). We defined its normal vector, pointing from the back to the front of the binding pocket, as the z axis. Along each ligand trajectory, we monitored both r and its z component (Figures 6C and S5B). The first increase of r to 9 Å was labeled as an unbinding event, whereas the next decrease of r to 1 Å was labeled as a rebinding event; and this label was repeated till the end of the trajectory. Depending on whether z was positive or negative when the unbinding or rebinding event occurred, the passage was through the front door or back door. Due to the large biases at the beginning of each simulation, the ligand very rapidly left the binding pocket. We started counting only from the subsequent rebinding event.

The total numbers of unbinding and rebinding events for each ligand are listed in Table 4. For the cognate ligands, unbinding shows a significant preference for the front door. Q_1_ has 14 events through the front door but only 4 events through the back door; for Q0, the counts are 11 versus 6. In contrast, the overwhelming preference for synthetic ligand unbinding is through the back door, with a total of 63 events. The total number of front-door unbinding events is only 9 for these ligands.

**Table 4.** Unbinding and rebinding events.

| Ligand | Unbinding |  | Rebinding |  |
| --- | --- | --- | --- | --- |
|  | Front | Back | Front | Back |
| $Q_1$ | 14 | 4 | 11 | 9 |
| $Q_0$ | 11 | 6 | 11 | 7 |
| $L_1$ | 5 | 17 | 7 | 16 |
| $L_2$ | 2 | 25 | 7 | 22 |
| $L_3$ | 2 | 21 | 3 | 23 |

The opposite preferences of the cognate and synthetic ligands also carry over to rebinding, though the preferences are somewhat blunted for both types of ligands. For cognate ligand rebinding, there is only a modest preference for the front door (11 events each for Q_1_ and Q_0_, compared with 9 and 7 events, respectively, for the back). For the synthetic ligand rebinding, the total number of events through the back door is 61, while the counterpart through the front door is 17. The preference for the back door is still significant but not as overwhelming as found for unbinding.

## DISCUSSION

We have investigated the molecular determinants underlying the binding of the PreQ_1_ riboswitch aptamer to cognate and synthetic ligands by combining the conventional MD simulations, free energy decomposition, and metadynamics simulations. The analyses on structural, energetic, and dynamic properties have advanced our understanding on both the overt differences between the cognate and synthetic groups of ligands and subtle differences within each group of ligands. In particular, the reduction of hydrogen bond donors and acceptors is the main reason for the decreased binding affinities of the synthetic ligands, while the increase in rings resulting in the opening of a back door to the binding pocket.

Our work has demonstrated the power of molecular dynamics simulations in completing structure determination and binding assays to provide crucial missing links. For example, the L2 loop is essential both for stabilizing ligands and for communicating ligand binding to the Shine-Dalgarno sequence for downstream signaling. Yet this loop is highly dynamic and nucleobases in the loop were cleaved (23) or subject to distortion by crystal contacts (25) in structure determination. Our MD simulations have now shown how this loop responds to the synthetic ligands (by extruding to form a base stack orthogonal to that formed when bound with the cognate ligands; Figure 4C) or to Mg^2+^ (which ties L2 to S1 to keep C15 in a position to form stable hydrogen bonds with the cognate ligands; Figure 3D). Moreover, our MD simulations have found that the methylamine head group of Q_1_ samples different torsion angles to alternate its hydrogen-bonding partner between G5 and G11, and L_3_ samples different poses to alternate its hydrogen-bonding partner between U6 and A29. Most interestingly, our MD simulations have revealed that ligands can enter and exit the binding pocket through multiple pathways and the cognate and synthetic ligands have opposite preferences. We hope that both our specific lessons on the PreQ_1_ riboswitch and our approach will provide guidance for designing riboswitch ligands in the future.

## DATA AVAILABILITY

Structures from molecular dynamics simulations and data files presented as plots in figures will be available, upon publication, on github at: https://github.com/hzhou43/PreQ1_Riboswitch

## FUNDING

This work was supported by National Institutes of Health Grant GM118091. GH’s participation in this work was also partially supported by Natural Science Foundation of Shandong Province Grant ZR2019MA040 and National Natural Science Foundation of China Grant 61671107.

## Notes

### Competing Interest Statement

The authors have declared no competing interest.

