## Supplementary Table and Figures for "Binding Free energy Decomposition and Multiple Unbinding Paths of Buried Ligands in a PreQ_1_ Riboswitch"

Short title: Multiple Unbinding Paths of Buried Ligands in a Riboswitch

Guodong Hu[1, 2] and Huan-Xiang Zhou[2, 3, *]

[1]Shandong Key Laboratory of Biophysics, Dezhou University, Dezhou 253023, China

[2]Department of Chemistry, University of Illinois at Chicago, Chicago, IL 60607

[3]Department of Physics, University of Illinois at Chicago, Chicago, IL 60607

*

*Supplementary Material*

**Table S1.** Binding free energies and their components (in kcal/mol) for five ligands in the absence $Mg^{2+}$.[a]

| | $Q_1$ | $Q_0$ | $L_1$ | $L_2$ | $L_3$ |
|---|---|---|---|---|---|
| $\Delta E_{\mathrm{ele}}$ | -24.34±2.24 | -16.44±1.98 | -6.07±0.79 | -4.98±0.42 | -6.3±0.41 |
| $\Delta E_{\mathrm{vdW}}$ | -33.67±0.29 | -35.52±0.22 | -41.68±0.64 | -47.25±0.67 | -36.38±2.11 |
| $\Delta G_{\mathrm{pol}}$ | 35.2±0.83 | 28.21±1.44 | 35.89±1.71 | 39.73±1.16 | 31.98±2.18 |
| $\Delta G_{\mathrm{nonpol}}$ | -3.32±0.01 | -3.24±0.00 | -4.2±0.03 | -4.7±0.08 | -3.94±0.11 |
| $\Delta E_{\mathrm{ele}} + \Delta G_{\mathrm{pol}}$ | 10.86±2.01 | 11.77±1.41 | 29.83±1.10 | 34.75±0.94 | 25.68±2.31 |
| $\Delta E_{\mathrm{vdW}} + \Delta G_{\mathrm{nonpol}}$ | -36.99±0.30 | -38.76±0.23 | -45.88±0.64 | -51.96±0.74 | -40.32±2.21 |
| $\Delta H$ | -26.13±1.73 | -26.99±1.35 | -16.06±0.75 | -17.2±1.05 | -14.64±0.56 |
| $T\Delta S$ | -17.29±0.14 | -18.58±0.23 | -16.26±0.55 | -15.09±0.67 | -17.37±0.43 |
| $\Delta G_{\mathrm{bind}}$ | -8.83±1.75 | -8.42±1.18 | 0.21±1.14 | -2.11±1.56 | 2.73±0.86 |

[a] These results were obtained from cMD simulations in the absence of $Mg^{2+}$. The terms have the same meanings as the counterparts in Table 2.

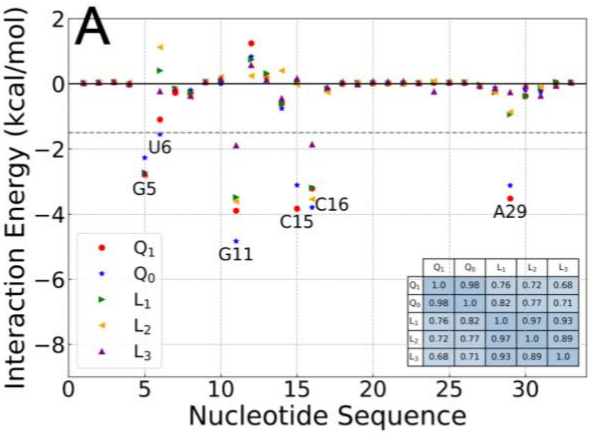


| | $Q_1$ | $Q_0$ | $L_1$ | $L_2$ | $L_3$ |
|---|---|---|---|---|---|
| $Q_1$ | 1.0 | 0.98 | 0.76 | 0.72 | 0.68 |
| $Q_0$ | 0.98 | 1.0 | 0.82 | 0.77 | 0.71 |
| $L_1$ | 0.76 | 0.82 | 1.0 | 0.97 | 0.93 |
| $L_2$ | 0.72 | 0.77 | 0.97 | 1.0 | 0.89 |
| $L_3$ | 0.68 | 0.71 | 0.93 | 0.89 | 1.0 |

**B**

| Ligand | G5 | | G11 | | C16 | |
|---|---|---|---|---|---|---|
| | In (%) | Dist (Å) | In (%) | Dist (Å) | In (%) | Dist (Å) |
| $Q_1$ | 47.68 | 3.53 | 99.91 | 3.50 | 99.60 | 3.46 |
| $Q_0$ | 42.37 | 3.54 | 100.00 | 3.46 | 100.00 | 3.46 |
| $L_1$ | 31.49 | 3.47 | 84.45 | 3.46 | 84.31 | 3.45 |
| $L_2$ | 42.25 | 3.46 | 99.90 | 3.42 | 99.82 | 3.41 |
| $L_3$ | 63.63 | 3.31 | 73.62 | 3.58 | 71.22 | 3.58 |

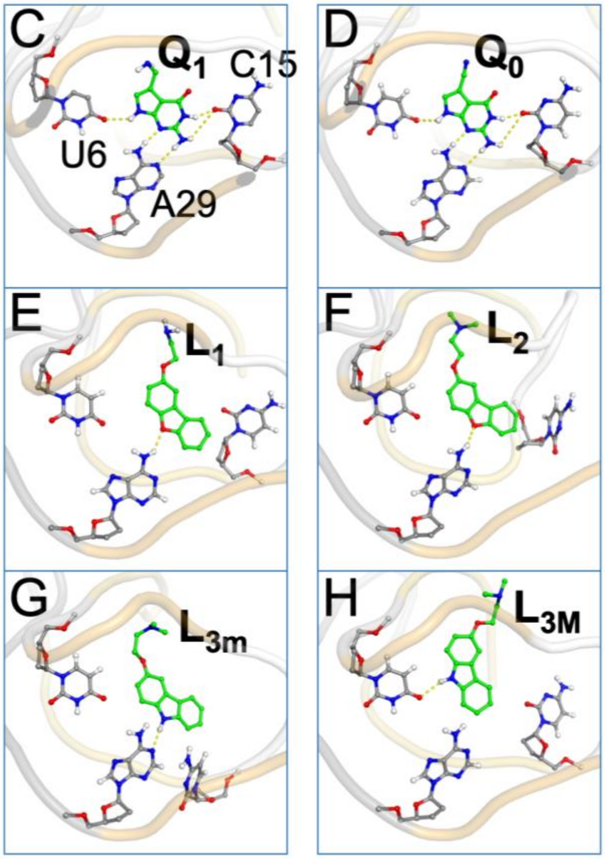


**Figure S1. Interactions of cognate and synthetic ligands with the PreQ$_1$ aptamer in cMD simulations without Mg$^{2+}$.** (A) Contributions of individual nucleotides to the binding free energies. A dashed horizontal line is drawn at -1.5 kcal/mol, which separates the pocket-lining nucleotides from the rest of the sequence. Inset: a table listing the correlation coefficients between the individual contributions of any two complexes. (B) In-fractions of three nucleobases and their average vertical distances from the ligand rings. (D)-(H) In-plane hydrogen bonds between ligands and nucleobases, shown as dashed lines, in representative conformations from cMD simulations.

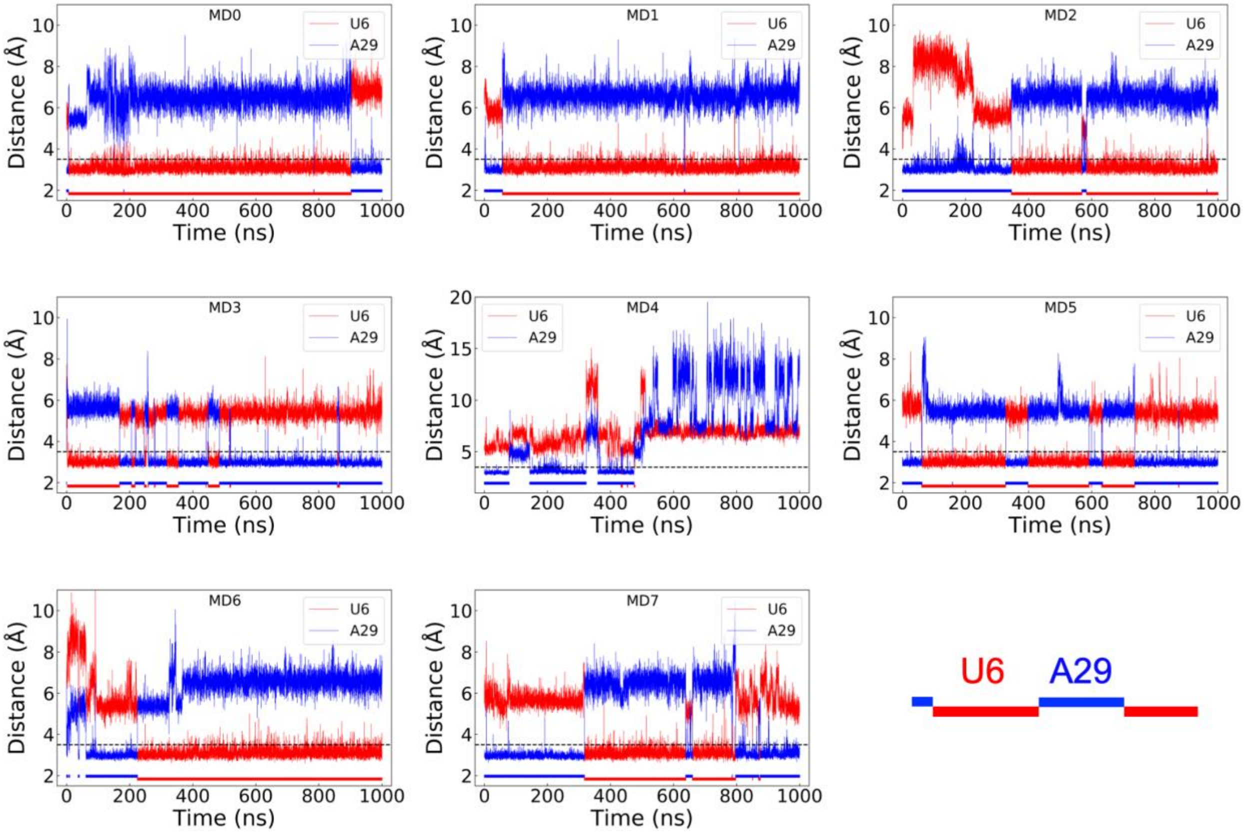


**Figure S2. Rapid switch of $L_3$ between two poses: one hydrogen-bonded with U6 and the other hydrogen-bonded with A29.** The distances of the $L_3$ N1 atom from the U6 O4 and A29 N1 atoms are shown as red and blue traces along the simulation time, in eight cMD simulations. In each panel, a horizontal dashed line is drawn at 3.5 Å. The horizontal bar at the bottom is colored red or blue, according to whether the U6 or the A29 distance is < 3.5 Å. The blue sections are raised slightly to better distinguish from the red sections. The bottom right panel shows an enlarged view of the blue and red sections of the horizontal bar. The MD4 simulation is special as the ligand partially slipped through the back door around 500 ns, pushing A29 out of the binding pocket; the ligand rings then flipped and retracked, leading to large distances from A29. Accordingly the upper bound of the ordinate is increased from 11 Å to 20 Å.

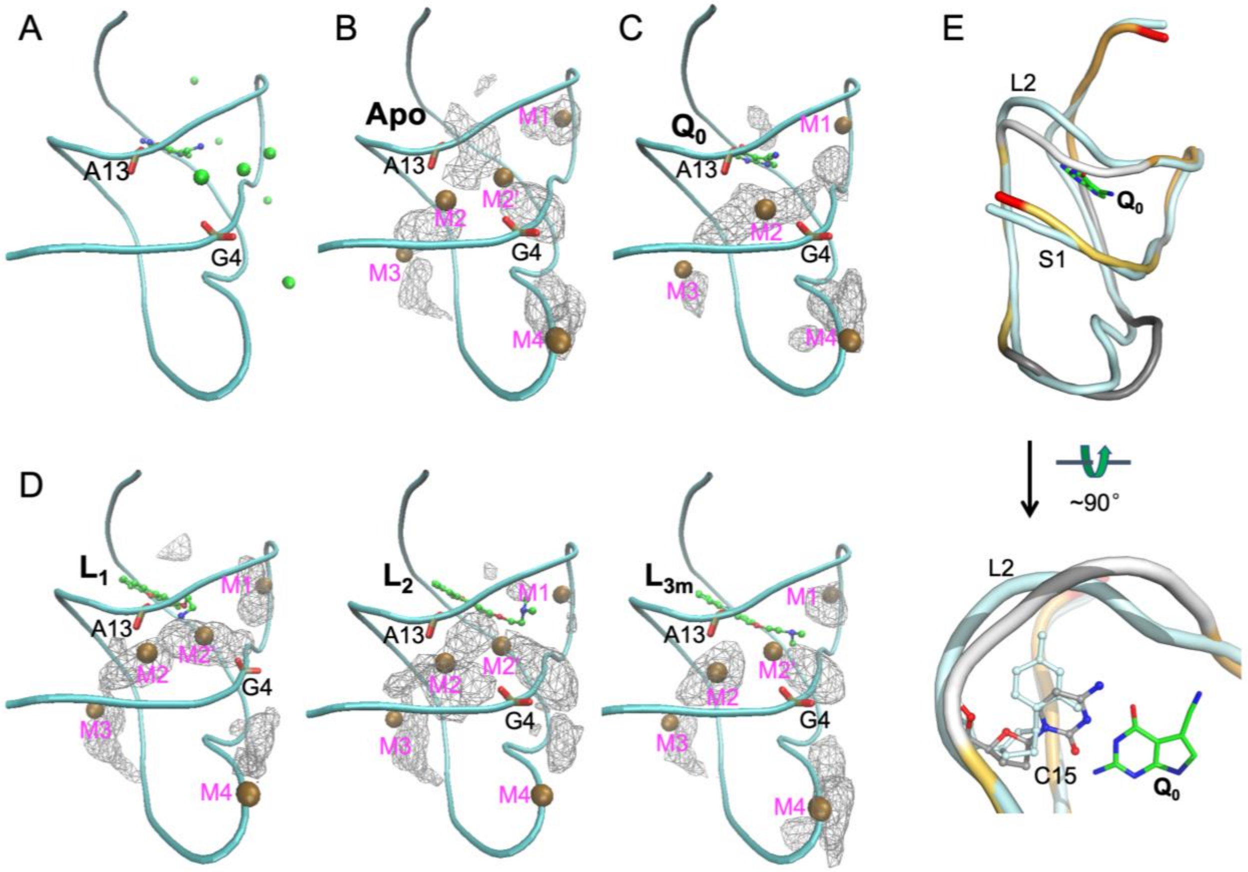


**Figure S3. Distributions and effects of $Mg^{2+}$.** (A) Seven $Mg^{2+}$ ions added by the MCTBI method, initially at shallow positions along two grooves of the aptamer. (B) Density contours of $Mg^{2+}$ ions in the apo form, shown as wireframe. Five $Mn^{2+}$ ions in PDB entry 6VUH are shown as ochre spheres; the corresponding $Mg^{2+}$ sites are labeled as M1, M2, M2', M3, and M4. Phosphate groups in G4 and A13 are shown in stick representation. (C) Corresponding presentation for the $Q_0$-bound form, except that four crystal $Mn^{2+}$ ions from PDB entry 6VUI are shown, with the sites labeled as M1, M2, M3, and M4. (E) Presentations for the $L_1$-, $L_2$-, and $L_{3m}$-bound forms, very similar to that shown in panel (B) for the apo form. (D) Effect of $Mg^{2+}$ ions on the separation of the L2 loop from the S1 helix in the $Q_0$-bound form. Two representative structures are superimposed, with the aptamer in the presence of $Mg^{2+}$ shown in the same multi-color scheme as in Figure 1A and the aptamer in the absence of $Mg^{2+}$ shown in a uniform cyan color. In the bottom view, the C15 nucleotides in the two structures are shown in a stick representation.

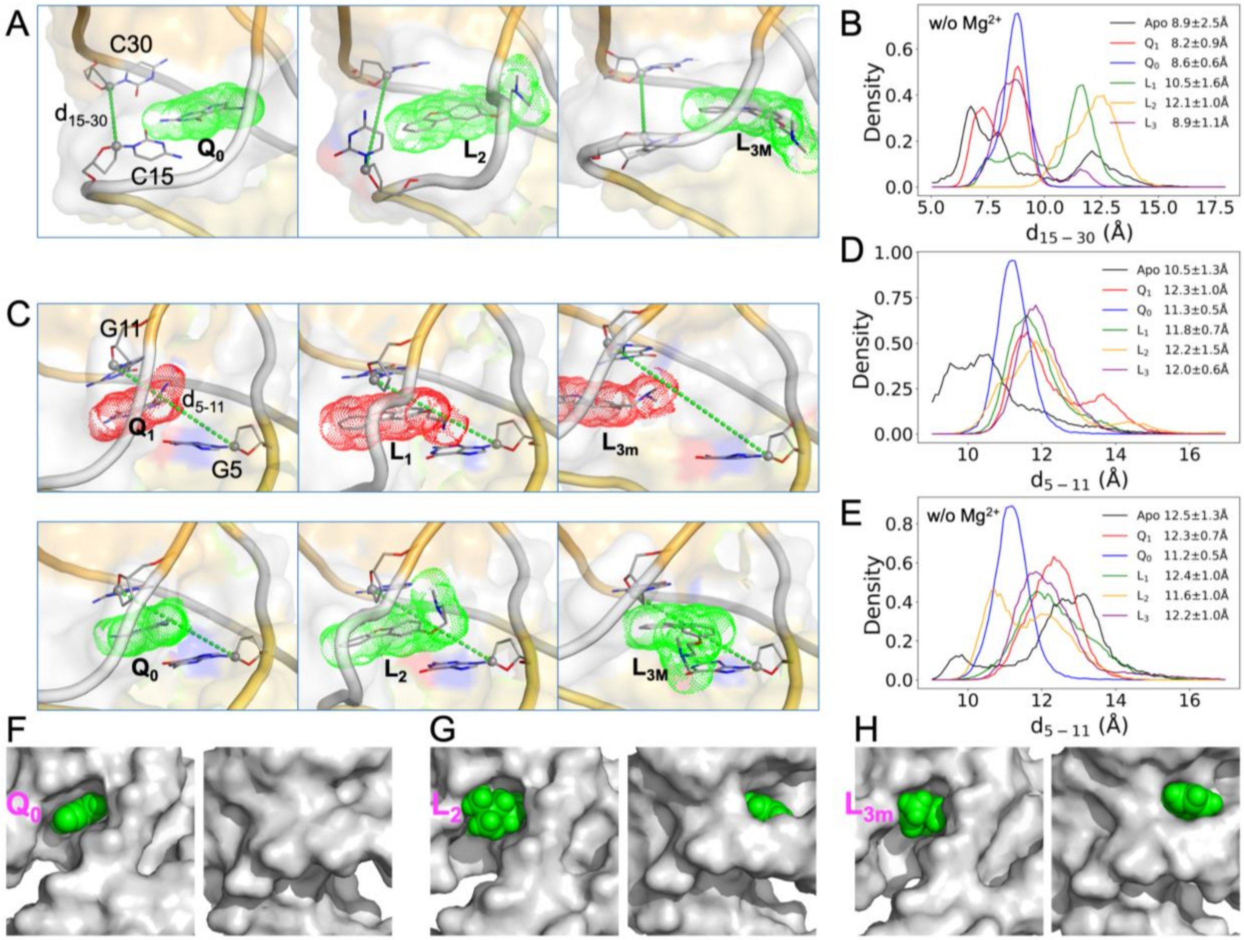


**Figure S4. Loosening in the back of the binding pocket when bound with synthetic ligands.** (A) Representative conformations of the $Q_0$-, $L_2$-, and $L_{3M}$-bound forms in cMD simulations with $Mg^{2+}$. Ligands are shown in both stick representation and as dot surface. The C1' atoms of C15 and C30 are connected to define the distance $d_{15\text{-}30}$. (B) The probability densities of $d_{15\text{-}30}$ in cMD simulations of the apo form and the five liganded forms without $Mg^{2+}$. (C) Representative conformations of the $Q_1$-, $Q_0$-, $L_1$-, $L_2$-, $L_{3m}$-, and $L_{3M}$-bound forms in cMD simulations with $Mg^{2+}$. The C1' atoms of G5 and G11 are connected to define the distance $d_{5\text{-}11}$. (D) The probability densities of $d_{5\text{-}11}$ in cMD simulations of the apo form and the five liganded forms with $Mg^{2+}$. (E) The probability densities of $d_{5\text{-}11}$ in cMD simulations of the apo form and the five liganded forms without $Mg^{2+}$. (F)-(H) Two views into $Q_0$, $L_2$, and $L_{3m}$, respectively, in the binding pocket. The aptamer is shown as gray surface while the ligands are shown as green spheres. The front and back views are shown on the left and right, respectively, in each panel.

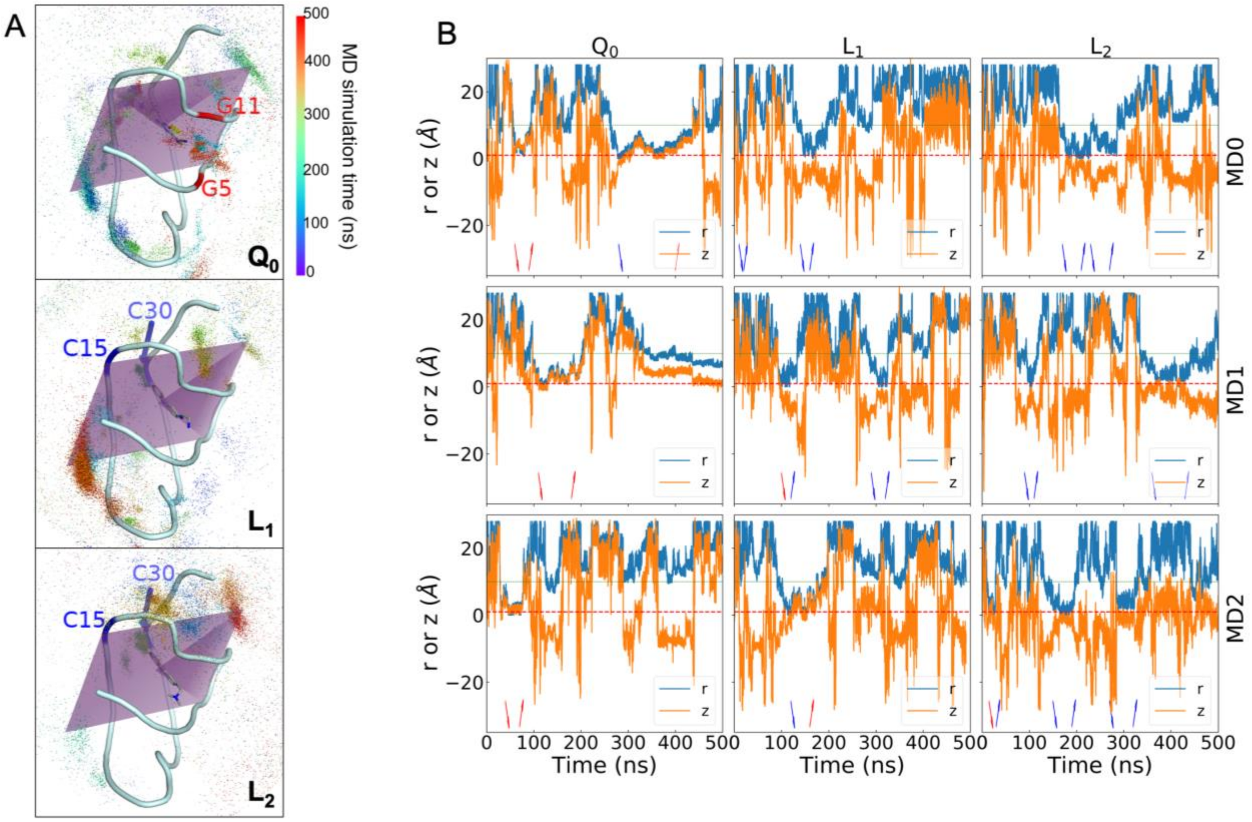


**Figure S5. Unbinding and rebinding pathways of ligands.** (A) The trajectories of ligand centers shown as dots colored according to the MD simulation time. The aptamer and bound ligands are shown in cartoon and stick representations, respectively. Top: $Q_0$; middle: $L_1$; and bottom: $L_2$. A plane in purple bisects the binding pocket into the front half and the back half. Two nucleotides defining the front door in the $Q_0$-bound complex are labeled in red; two nucleotides defining the back door in the $L_1$- and $L_2$-bound complexes are labeled in blue. (B) Time traces of r and z in three metadynamics simulations.
